# Methods for estimation of model accuracy in CASP12

**DOI:** 10.1101/143925

**Authors:** Arne Elofsson, Keehyoung Joo, Chen Keasar, Jooyoung Lee, Ali H. A. Maghrabi, Balachandran Manavalan, Liam J. McGuffin, David Ménendez Hurtado, Claudio Mirabello, Robert Pilstål, Tomer Sidi, Karolis Uziela, Björn Wallner

**Affiliations:** Department of Biochemistry and Biophysics and Science for Life Laboratory, Stockholm University, Box 1031, 171 21 Solna, Sweden; Department of Physics, Chemistry, and Biology, Bioinformatics Division, Linköping University, 581 83 Linköping, Sweden; School of Biological Sciences, University of Reading, Whiteknights, Reading, RG6 6AS, United Kingdom; Department of Computer Science, Ben Gurion University of the Negev, Israel; Center for In Silico Protein Science and Center for Advanced Computation, Korea Institute for Advanced Study, Seoul 130-722, Korea; Center for In Silico Protein Science and School of Computational Sciences, Korea Institute for Advanced Study, Seoul 130-722, Korea

**Keywords:** protein structure prediction, quality assessment, CASP, Estimates of model accuracy, Consensus predictions, Machine learning

## Abstract

Methods for reliably estimating the quality of 3D models of proteins are essential drivers for the wide adoption and serious acceptance of protein structure predictions by life scientists. In this paper, the most successful groups in CASP12 describe their latest methods for Estimates of Model Accuracy (EMA). We show that pure single model accuracy estimation methods has shown clear progress since CASP11; the three top methods (MESHI, ProQ3, SVMQA) all perform better than the top method of CASP11 (ProQ2). The pure single model accuracy estimation methods outperform quasi-single (ModFOLD6 variations) and consensus methods (Pcons, ModFOLDclust2, Pcomb-domain and Wallner) in model selection, but are still not as good as those methods in absolute model quality estimation and predictions of local quality. Finally, we show that when using contact based model quality measures (CAD, 1DDT) the single model quality methods perform relatively better.

## Introduction

Estimates of Model Accuracy (EMA) have been apart of protein structure prediction since its infancy. It is actually built into virtually all methods as the energy functions that they optimize. Yet, these energy functions provide only relative accuracy estimate, with moderate power in properly ranking models. Further, when one tries to use models from different methods, their associated energies are not comparable. Thus, accurate posterior quality estimation methods are essential for protein structure prediction to fulfill its potential.

Motivated by the intriguing experiment of Novotny et al.^11^ early model accuracy assessment methods focused on distinguishing wrong models (or decoys) from the native structure^2,3^. Knowledge based energy functions were developed to solve this problem and to guide protein folding and fragment assembly simulations and for threading studies. Notably, the methods by Sippl, which used a knowledge based energy function for threading, were quite successful in CASP1-3^4,5^. However, in later CASP experiments threading methods have not been able to keep up with methods that use evolutionary information from the rapidly growing sequence databases.

None of the energy functions that were developed to distinguish native and non-native protein models showed any major success in CASP. Instead, more successful methods, starting with ProQ^6^, that aim to predict the quality of a model have been more successful. One of the notable features separating these methods from the earlier knowledge based energy terms were the use of compatibility with predicted structural features, such as secondary structure. These methods are nowadays referred to as single model quality assessment methods to distinguish them from methods that use clustering (or consensus) of many models. Since the introduction of ProQ other methods based on the same idea has been introduced, including QMEAN^7^ that has performed on par with ProQ in earlier CASPs. In earlier CASPs the single methods have not been as successful as the methods that take into account structural similarity of models, i.e. consensus based methods^8^, but since CASP11 they perform at least on par with the consensus methods in at least some of the tasks^8^.

The first successful attempt of independent model quality estimation, in the context of CASP, was when the first meta predictor was introduced in CASP4^9^, where it was shown that combining the results from several servers could provide better models than any of the individual servers. However, in CASP4 the model quality estimates were done manually. From this exercise it was realized that a simple rule combining the predictions from several servers could outperform all individual servers. This algorithm chose the most frequent fold predicted by all servers, i.e. choosing the consensus^9,10^.

Soon after CASP4, the first automatic consensus method, Pcons, was introduced^11^. This was later followed by a simpler (and more robust) method, 3D-Jury^12^. Later versions of Pcons are very similar to 3D-Jury^13^, the only difference is in the details of the superposition method. In CASP5 it was clear that these methods could be used to outshine all individual servers if the results were combined. In CASP7 model accuracy estimation became a category by itself for the first time ^1414^.

Quasi-single model methods, such as the latest ModFOLD servers^15,16^ compare a model with models generated by a local prediction-pipeline using the consensus approach. These methods, as well as Pcomb^13^ that uses the Pcons consensus approach, combine the consensus score with one or several pure single model approaches. The performance of the best quasi-single approaches often match the performance of the consensus methods, but with the ability to evaluate a single model at a time given that a set of external predictions exist.

Below, we will first describe shortly the methods used by our groups in CASP12. Thereafter, we will compare their performance and discuss our insights about their pros and cons.

## Methods

A summary of all methods discussed in this paper is presented in Table 1. Below, each group presents their methods briefly.

**Table 1.** Summary of the best performing QA methods in CASP 12 and comments about their strength and weaknesses. Methods basically identical have been merged

| Methods | Type | Comment about Global performance | Comment about Local Performance |
| --- | --- | --- | --- |
| MESHI<sup>36</sup> | Single | Top model selection | N/A |
| MESHI_con<sup>36</sup> | Single* | Top model selection | N/A |
| ProQ2<sup>21</sup> | Single | Good model selection | Acceptable local scores |
| ProQ3<sup>17</sup> | Single | Top model selection | Good local scores |
| SVMQA<sup>45</sup> | Single | Top model selection | N/A |
| ModFOLD6<sup>16</sup> | Quasi-single | Balanced performance | Good assignment of local scores |
| ModFOLD6_rank<sup>16</sup> | Quasi-single | Acceptable model selection | Identical to ModFOLD6 |
| ModFOLD6_cor<sup>16</sup> | Quasi-single | Best absolute but suboptimal model selection | Identical to ModFOLD6 |
| qSVMQA<sup>45</sup> | Quasi-single | Assignment of the absolute score is not accurate. | N/A |
| ModFOLDclust2<sup>25</sup> | Clustering | Good assignment of absolute global scores but suboptimal model selection | Top assignment of local scores |
| Pcons<sup>11</sup> | Clustering | Good assignment of absolute global scores | Top assignment of local scores |
| Pcomb-domain<sup>13</sup> | Combined | Good assignment of absolute global scores, requires good domain prediction | Top assignment of local scores |
| Wallner | Combined | Good assignment of absolute global scores | Top assignment of local scores |

\* = MESHI\_con is not pure single methods but requires multiple models to average the predictions

### Elofsson group

We participated with several accuracy estimation methods in CASP12. Here, we will highlight the two methods that performed best; the single model accuracy estimation tool ProQ3^17^ and our consensus based method Pcons^11^. Our other methods included an early version of ProQ3D^18^ the deep learning version of ProQ3. ProQ3_diso is a version of ProQ3 where disordered residues are ignored and RSA_SS is a simple quality assessment method that only utilizes predicted secondary structure and surface area. For details see the CASP12 abstracts at http://predictioncenter.org/casp12/doc/CASP12_Abstracts.pdf.

**ProQ3^17^** is the latest version of our single model accuracy estimation methods^6,19–21^. In Table 2 we describe the most important developments in the history of ProQ. In addition to using the same descriptions of a model as ProQ2^21^ it also uses Rosetta energy functions. All input features are combined together to train a linear SVM. The training data set is a subset of CASP9 with 30 models per target. We also tested a few developmental methods of ProQ in CASP12, but none of these performed significantly better than ProQ3 and are therefore not discussed here. However, it can be noted that we have recently developed an improved version of ProQ3, ProQ3D that uses a deep-learning approach but identical inputs as ProQ3^18^. The final version was not ready for CASP12 and the preliminary version used did not perform better than ProQ3. ProQ3 is available both as source code from https://bitbucket.org/ElofssonLab/proq3, and as a web-server at http://proq3.bioinfo.se/.

**Table 2.** Description of the evolution of ProQ methods used in CASP and their relative performance.

| Method | Major Novelty | Correlation global/local |
| --- | --- | --- |
| **ProQ**<sup>6</sup> | First method trained to predict “quality” of a model. Using a combination of structural descriptions and agreement with predicted secondary structure. | 0.71<sup>A</sup>/- |
| **ProQres**<sup>20</sup> | Predicting *local* qualities - global quality is sum of local quality. | -/0.56<sup>B</sup> |
| **ProQ2**<sup>21</sup> | Global agreement with predicted RSA and SS plus profile weighting. Uses a linear kernel SVM. | 0.80/0.71<sup>B</sup><br>0.84/0.72<sup>C</sup> |
| **ProQ3**<sup>17</sup> | Added rosetta energies to the inputs. | 0.87/0.74<sup>C</sup> |
| **ProQ3D**<sup>18</sup> | Linear kernel SVM is replaced by a two-layer perceptron. | 0.91/0.77<sup>C</sup> |

<sup>A</sup> from original ProQ publication <sup>6</sup>
<sup>B</sup> from ProQ2 publication <sup>21</sup>
<sup>C</sup> on CASP11 dataset trained on CASP9 and CASP10

**Pcons^11^** is used with default setting. This means that the score is calculated by performing a structural superposition using the algorithm described by Levitt and Gerstein^22^ of a model against all other models. To avoid bias, comparisons between models from the same method are ignored. After superposition, the “S-score” is calculated for each residue in the model^23^. The average S-score for all residues and pairs of models is then used to calculate the final Pcons score. For local predictions, the average S-score is converted to a distance as described before^13^. Pcons is freely available from https://github.com/biomwallner/Pcons/. It should be noted that a number of heuristic optimizations have been implemented in Pcons to enable the pairwise comparison of hundreds of proteins in a short time ^24^.

### McGuffin Group

We participated in CASP12 with three new quasi-single model method variants, ModFOLD6, ModFOLD6_cor and ModFOLD6_rank (Figure 1), and one older clustering method, ModFOLDclust2.

**Figure 1:**
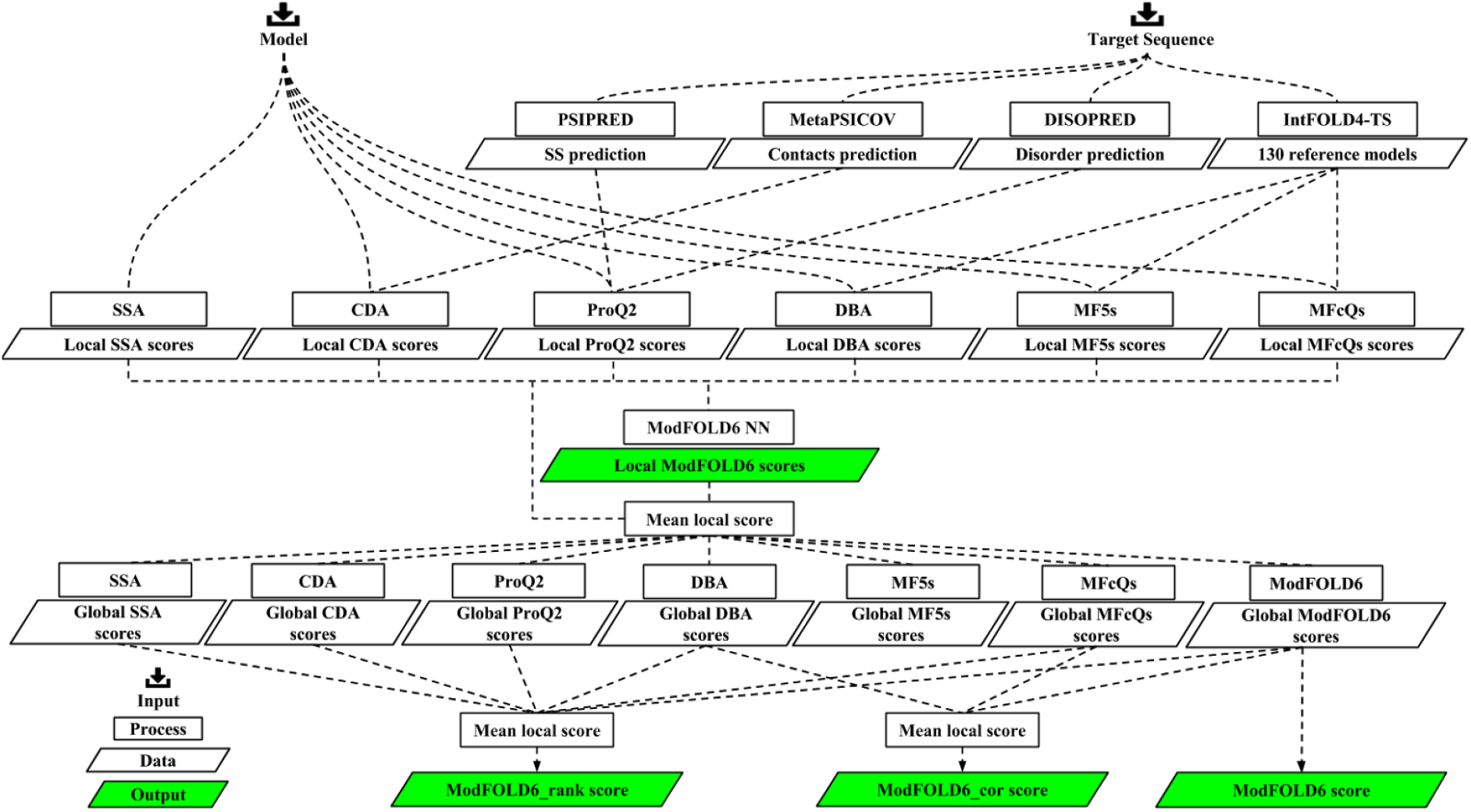
Flowchart outlining the principal stages of the ModFOLD6 server prediction pipeline. The initial input data are the target sequence and a single 3D model. The output data are the local/per-residue scores from the ModFOLD6 NN and the global score variants - ModFOLD6, ModFOLD6_rank and ModFOLD6_cor. The ModFOLD6 pipeline is dependent on the following methods PSIPRED^42^, DISOPRED^53^ and MetaPSICOV^54^.

#### ModFOLD6

The ModFOLD6 server^16^ is the latest version of our freely available public resource for the accuracy estimation of 3D models of proteins^15,25,26^. The ModFOLD6 server combines a pure-single and quasi-single model strategy for improving accuracy of local and global model accuracy estimates. Our initial motivation in the development of ModFOLD6 was to increase the accuracy of local/per-residue assessments for single models^16^.

For the local/per-residue error estimates, each model was considered individually using two new pure-single model methods, the Contact Distance Agreement (CDA) and the Secondary Structure Agreement (SSA) scores^16^, as well as the best pure single method in earlier CASPs, ProQ2^21,27^. Additionally, three alternative quasi-single model methods were used to score models including: the newly developed Disorder B-factor Agreement (DBA), the ModFOLD5_single (MF5s) and the ModFOLDclustQ_single scores (MFcQs)^16^ - each of which made use of a set of 130 reference 3D models that were generated using the latest version of the IntFOLD-TS^28,29^ pipeline from the IntFOLD server^30,31^. The component per-residue scores from each of the 6 alternative scoring methods, mentioned above, were combined into a single score for each residue using an Artificial Neural Network, which was trained to learn the local S-score^23^ as the target function^16^ (i.e. the same target function as ProQ2, described below and in Table 2 was used, but with d_0_ set to 3.9).

For global scoring, in the ModFOLD6 variant we simply took the mean local score for each model (i.e. the sum of the per-residues scores divided by the target sequence length). However in our internal benchmarks, using CASP11^8^ and CAMEO^32^ data prior to CASP12, we realized that simply taking the mean per-residue score from ModFOLD6 alone was not optimal and performance differed depending on the intended use case, i.e. selecting the best models or accurately reproducing the model-target similarity scores. Therefore we also exhaustively explored all linear combinations of each of the alternative global scores, in order to find the optimal mean score (OMS) for each major use case^16^.

#### ModFOLD6_cor

The aim of developing the ModFOLD_cor global score variant was to optimize the correlations of predicted and observed global scores i.e. the predicted global accuracy estimation scores produced by the method should be close to linear correlations with the observed global accuracy estimation scores. The OMS for the ModFOLD6_cor global score was found as:

ModFOLDclustQ_single_global + DBA_global + ModFOLD6_global)/3

where the _global suffix indicates that the mean local score was taken for the scoring method indicated above.

#### ModFOLD6_rank

The aim of developing the ModFOLD6_rank global score variant was to optimise for the selection of the best models i.e. the top ranked models (top 1) should be closer to the highest accuracy, regardless of the relationship between the absolute values of predicted and observed scores. The OMS for the ModFOLD6_rank global score was found as:

ModFOLD6_rank=ModFOLDclustQ_singlc_global + ProQ2_global + CDA_global + DBA_global + SSA_global + ModFOLD6_global)/6.

Note that the local scores submitted for each of the three ModFOLD6 variants were identical and it was only the global scores (and therefore the ranking of models), which differed between the three ModFOLD6 variants. All three of the ModFOLD6 variants are freely available at: http://www.reading.ac.uk/bioinf/ModFOLD/ModFOLD6form.html

#### ModFOLDclust2

The ModFOLDclust2 method^33^ is a leading automatic clustering based approach for both local and global 3D model accuracy estimation assessment^8,34,35^. The ModFOLDclust2 server tested during CASP12 was identical to that tested during the CASP9, CASP10 & CASP11 experiments. The local and global scores have been previously described^33^ and are unchanged since CASP9. Thus, the ModFOLDclust2 method serves as a useful gold standard/benchmark against which progress in the development of single model methods may be measured. ModFOLDclust2 can be run as an option via the older ModFOLD3 server (http://www.reading.ac.uk/bioinf/ModFOLD/ModFOLD_form_3_O.html). The ModFOLDclust2 software is also available to download as a standalone program (http://www.reading.ac.uk/bioinf/downloads/).

### Keasar Group

We participated in CASP12 with two EMA methods, MESHI-score (implemented by the MESHI_server group) and MESHI-score-con (MESHI_con_server), the latter is a slight variation on the former. Below we first present the general scheme, which is used by both methods, and then conclude with the variations tried in MESHI-score-con.

While preliminary versions of MESHI-score were used in CASP10 and CASP11, it has reached stability only after CASP11^36^. The software architecture, however, is modular, extendable by design, and under continuous development. Thus, the version that took part in CASP12 was more advanced than the one presented earlier^36^.

The MESHI-score pipeline (Figure 2) starts with a regularization step that includes sidechain repacking by SCWRL^37,38^ and restrained energy minimization (Figure 2 *II*). This step sharpens the quality signal of structural features by reducing noise, which is due to peculiarities of decoy generating methods. Features are extracted from the regularized structures (Figure 2 *III*) and fed to an ensemble of 1000 independently trained predictors (Figure 2 *IV*). Each predictor outputs (*S_i_, W_i_*), a pair of an EMA score and weight (Figure 2 *VI*). The weighted median of this set of pairs is the final MESHI-score (Figure 2 *VII*). In addition, we also calculate the weighted interdecile range and entropy of the pairs set:

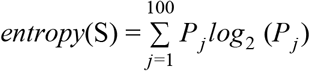

where S is the set of 1000 (*s_i_w_i_*) pairs and

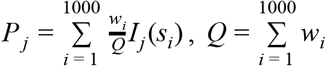

And

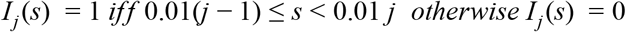

**Figure 2:**
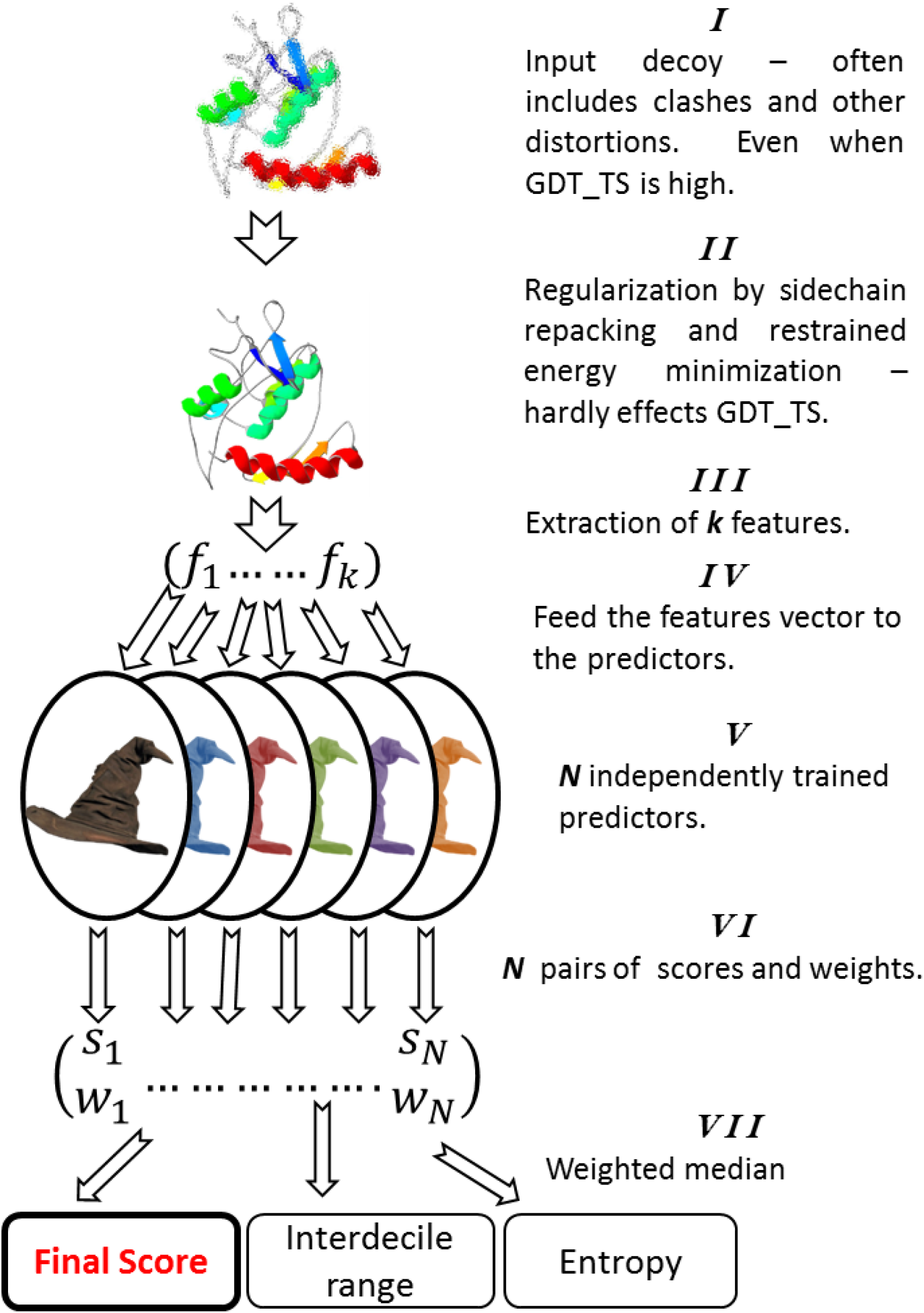
The MESHI-score pipeline starts with a regularization step that includes sidechain repacking by SCWRL^37,38^ and restrained energy minimization. Features are extracted from the regularized structures and fed to an ensemble of independently trained predictors. Each predictor outputs a pair of values: an EMA score and weight, and the weighted median of this set of pairs is the final MESHI-score.

The larger these numbers are the less reliable is the score, as they suggest disagreement between the predictors.

The feature set that was used in CASP12 included 82 features (for details see https://www.cs.bgu.ac.il/~frankel/TechnicalReports/2015/15-06.pdf)

These features may be clustered into nine broad categories:

1. Pairwise energy terms, which represent interactions between atoms, adopted from the literature^39–41,39–41^.
2. Compatibility of the decoy secondary structure and solvent accessibility with their PSIPRED^42^ prediction.
3. Standard bonded energy terms (e.g., quadratic bond term).
4. Torsion angle terms (compatibility with Ramachandran plot and rotamer preferences^43^)
5. Hydrogen bond terms^44^
6. Solvation and atom environment terms that quantify the cooperativity between hydrogen bond formation and atom burial.
7. Radius of gyration, and contact terms that quantify the compatibility of decoys with the expected, length dependent, ratios between the radii of gyration and numbers of contacts in different subsets of protein atoms (e.g., polar and hydrophobic).
8. Meta-features that quantify the frustration within decoys (native structures tend to be minimally frustrated) by considering the distribution of the pairwise and torsion energies within the decoys.
9. Combinations of the above features, which were developed in previous studies ^36^.

The predictors (Figure 2 *V*) are nonlinear functions that get feature vectors as an input and output a pair of numbers: an EMA score, and a weight that represents the reliability of the score. The parameters of the predictor functions, as well as the subset of features that they use, are learned by stochastic optimization. Each predictor is trained to minimize a different objective function and thus tends to be more sensitive in a specific GDT_TS subrange. Scores within the predictor’s sensitivity region are considered more reliable and thus, have a higher weight. A more detailed description of the predictor’s training may be found in Mirzaei et al^36^.

MESHI-score-con is a variant on the MESHI-score theme, which aims to improve the consistency MESHI-score by a post processing step that takes into account the similarities between decoys. Ideally, after regularization (Figure 2 *II*) very similar decoys should produce similar feature vectors, and thus have similar MESHI-scores. Yet, careful examination of MESHI-score results indicates that this is not always the case, and often very similar decoys have quite different scores. MESHI-score-con aims to alleviate this problem by improving the agreement between the scores of very similar decoys. To this end, we associate the MESHI-score of each decoy with a weight, which is inversely proportional to the entropy of the score-weight pairs (Figure 2 *VII*). We also associate each decoy with a neighbors-set that includes very similar (GDT_TS >= 95) neighbors as well as the decoy itself. MESHI-score-con is a weighted average of the decoy’s MESHI-score and the average score of its neighbor-set. Thus, a low weight decoy (presumably a less reliable one) with higher weight neighbors is strongly biased towards the average score of its neighbors. Yet the score of a decoy without neighbors is unaffected regardless of its weight. Thus, unlike consensus methods MESHI-score-con may pick an exceptionally good decoy.

### Lee Group

We participated in CASP12 with two methods, namely SVMQA and quasi-SVMQA (qSVMQA). qSVMQA augments TM-score between GOAL_TS1 and the server model with an appropriate value of weight *w* to the SVMQA score:

qSVMQA = SVMQA + *w* * (TM-score between GOAL_TS1 and the server model).

The value of *w* was set separately for stage1 models (0.84) and for stage2 models (0.15). We determined the optimal value of *w* using CASP11 single-domain targets. Below, we briefly describe SVMQA and highlight its results in the model selection of stage2 targets in CASP12.

SVMQA is a support-vector-machine-based protein single-model global QA method. SVMQA predicts the global QA score as the average of the predicted TM-score and GDT_TS score by combining two separate predictors, SVMQA_GDT and SVMQA_TM. For SVMQA we used 19 features (8 potential energy-based terms and 11 consistency-based terms between the predicted and actual values of the model) for predicting the QA score (TM-score or GDT_TS score). Among these 19 features, 3 features (orientation dependent energy, GOAP angular energy and solvent accessibility consistency score) were not used in earlier versions, while the other 16 have been used in existing methods. The description of each feature along with the selection of the final set of SVM parameters and the final set of features for these two predictors have been published recently^45^. In short, SVMQA_TM uses all of the 19 features to predict TM-score of a given model, whereas SVMQA_GDT uses only 15 features to predict the GDT_TS score.

In CASP11, we used our old QA method, RFMQA^46^. The result of RFMQA on CASP11 targets was quite successful but not as good as that of SVMQA on CASP12 targets. Prior to CASP12, we benchmarked the performance of SVMQA on CASP11 targets and compared it to that of RFMQA, and we found that SVMQA significantly outperformed RFMQA in terms of both ranking models and selecting a more native-like model. The major updates of SVMQA over RFMQA is as follow: (i) The choice of machine learning method was different, an SVM (support vector machine) was used in SVMQA while a random forest was used in RFMQA; (ii) we used CASP8-9 domain targets as the training dataset for RFMQA, while CASP8-10 domain targets were used in SVMQA; (iii) 19 input features were used in SVMQA, whereas, only 9 of these features were used in RFMQA; (iv) The objective function to train for RFMQA was TM_loss_ (difference between the TM-score of the selected model and the best TM-score), while that for SVMQA was the correlation coefficient between the actual ranking and the predicted ranking; and (v) SVMQA used two separate predictors for TM-score and GDT_TS score, while RFMQA used only a predictor for TM-score.

### Wallner group

We participated with three EMA methods; ProQ2^21^, Pcomb-domain, and Wallner.

**ProQ2** is a single model accuracy estimation program based on a linear kernel support vector machine trained on a set of structural descriptors of a model. ProQ2 is trained to predict the local S-score^23^:

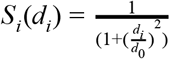

where d_i_ is the local distance deviation for residue i in the optimal superposition that maximize sum of S over the whole protein, and d_0_ is a distance threshold put to 3.0 here. The global score is the sum of local S_i_ divided by the target length yielding a score in the range [0,1]. Local S-scores, S_i_, were converted to local distance deviation using the formula:

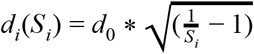

ProQ2 has participated in CASP since CASP10. Before CASP11 we implemented ProQ2 as a scoring function in Rosetta^27^, enabling scoring and integration in any Rosetta protocol. ProQ2 was top-ranked in both CASP10 and CASP11. This inspired developers of novel methods including SVMQA, MESHI-score and ProQ3. ProQ2 is further included in several hybrid methods incorporating ProQ2 directly for improved model accuracy estimation, laying the foundation for the improvement we see in some of the top-ranked methods in the current CASP12, e.g. Wallner, Pcomb, ModFOLD6^16^ in QA predictions, and the BAKERROSETTA-SERVER^47^ and the IntFOLD4 server for TS prediction.

**Wallner** method in this CASP is what was called Pcomb in earlier CASPs^13,48^ that combines ProQ2 and Pcons using the linear combination:

Pcomb=0.2 * ProQ2+0.8*Pcons

for global prediction^20^. For local prediction the same formula was used to calculate weighted local S-scores, which then were converted to distances using the d_j_(S_j_) formula, described above.

**Pcomb-domain** method is a new domain-based version of Pcomb. Traditionally, consensus methods, including Pcons^1111^ (https://github.com/bjomwallner/pcons), have always used rigid-body superposition for the full-length models, thereby selecting models that overall have the highest consensus, ignoring the fact that smaller domains from other models could have higher consensus over that region. To try to overcome this problem we developed a domain-based version of Pcons, which uses an initial domain definition. The domain-based Pcons scores were combined with local predicted scores from ProQ2. Two different methods were used to predict the domain boundaries of the target sequences, the first used the domain definitions from the Robetta server and the second was based on spectral analysis of the top ranking server models according to the regular Pcomb method. The results from these two methods were manually evaluated to decide the final domain boundaries. In addition, the Pcons and ProQ2 scores were weighted in a slightly different way compared the regular Pcomb method; following a parameter optimization based on targets released in the last two editions of CASP the relative weight for ProQ2 was increased to 0.3 resulting in this formula:

Pcomb-domain=0.3 * ProQ2-domain+0.7* Pcons-domain

Furthermore, d_0_ was increased from 3.0Å to 5.0Å as it showed improved model selection on CASP11 data, increasing the d_0_ shifts the sensitivity to detect differences for lower quality (higher RMSD) residues. As for both ProQ2 and Pcons all predictions are performed in the S-score space, global scores are sum of local scores, and the local S scores are transformed to distances in the final step, using the d_i_(S_i_) formula above.

### Results

A detailed analysis of CASP12 EMA methods is provided in the accompanying EMA assessment paper^49^. In this section, we refer to the results provided in this paper pertaining to our methods and also perform an additional analysis based on the correlation between different scores for different types of methods.

#### Global accuracy estimations in CASP12

In the EMA assessment paper^49^ the accuracy of CASP12 methods in selecting the best model according to the GDT_TS score is shown. Three single model accuracy estimation methods are ranked at the top in terms of identifying the best model with the average error (i.e., difference between the GDT_TS of the selected model and the best GDT_TS) around 5 GDT_TS units. The individual ranking of these methods depends on the evaluation criteria and according to the assessment paper^49^ the difference between the top methods is not significant. The best consensus and quasi-single methods are only slightly worse than the pure single methods using these criteria. However, this is a significant progress since last CASP.

In the Figure 5 of the accompanying paper^49^ the ability to distinguish between good and bad models it is clear that the best methods use consensus or quasi-single methods and combine them with single model approaches, at least when using GDT_TS for evaluation. The top three methods are using the single model method ProQ2 as part of the their scoring. Wallner and Pcomb-domain scores are weighted sums of ProQ2 and Pcons scores, while ModFOLD6_rank uses ProQ2 together with many other scores. Still, even though the top methods are statistically better^49^, the much simpler pure consensus methods Pcons and ModFOLDclust2 are not far behind ranked 6th and 9th when using S-score and even better than ModFOLD6 when using 1DDT.

The ability of methods to rank the top models for each target was evaluated using the per target correlation, i.e. the correlation of estimated and observed accuracy for each target. In Figure 3, the distribution of per target correlation for the methods studied here and the three different model accuracy estimation measures are shown. The distributions are sorted by the median. It can be seen that the individual rankings of the methods are quite different depending on which accuracy measure that is used. When using GDT_TS^50^, consensus and quasi-single based methods clearly outperform the single model accuracy estimation methods. In contrast when using CAD^51^ or 1DDT^52^ the best correlation is obtained with ProQ3 and all the top methods are single model accuracy estimations. A similar difference in ranking can be seen in the AUC analysis on the CASP homepage (http://predictioncenter.org/caspl2/qa_aucmcc.cgi). Here, ProQ3 is ranked 20th when using GDT_TS but 7th when using CAD. In contrast Pcons is ranked as 4th using GDT_TS and as 12th using CAD. Interestingly, it can be seen that the “pure” consensus methods (Pcons, MODFOLDClust2) show only a modest per target correlation with CAD or 1DDT, see Figure 3.

**Figure 3:**
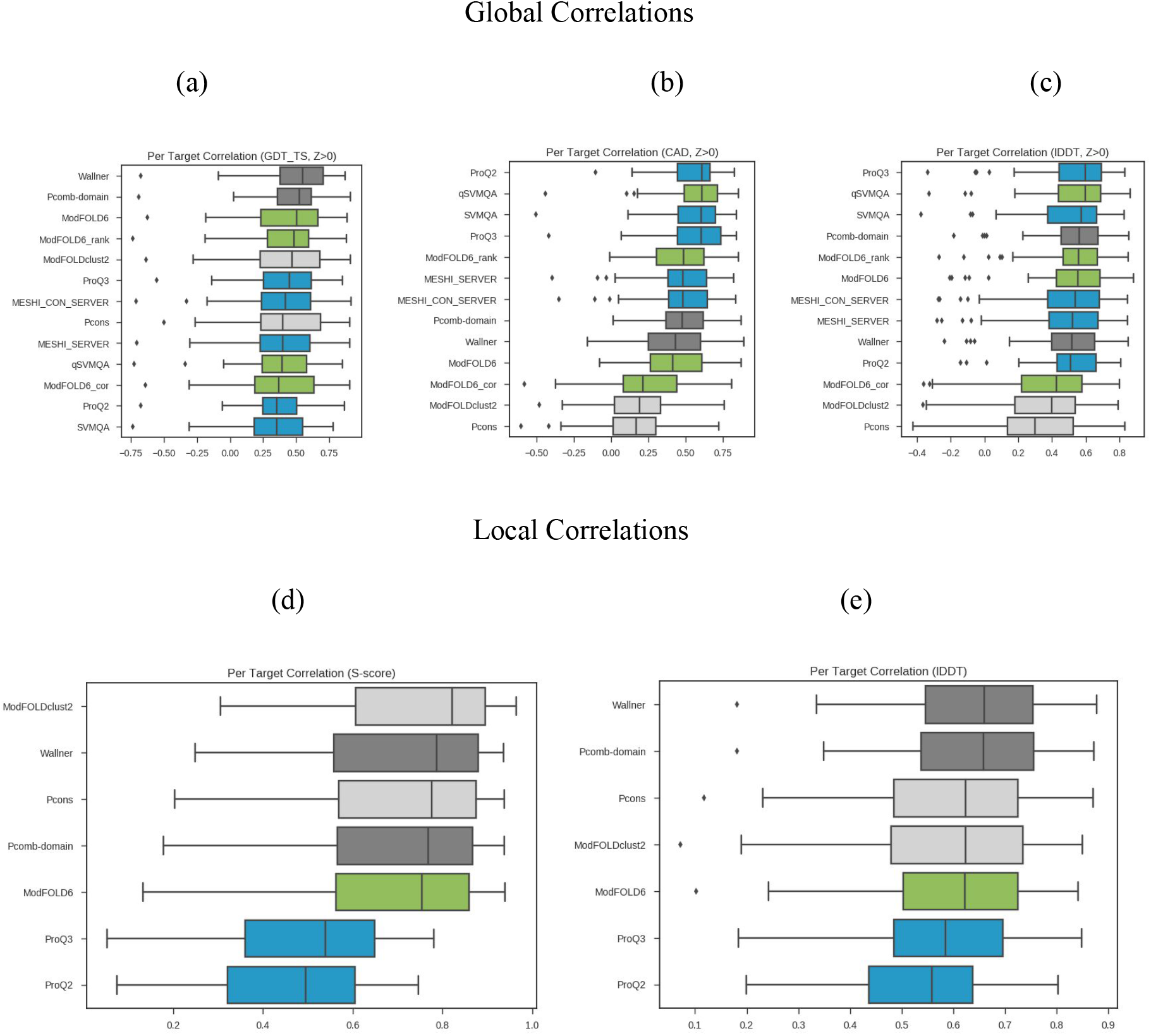
Boxplots of per target correlation for the methods presented in this paper for GDT_TS, CAD, and 1DDT, (a)-(c) global evaluations, (d)-(e) local evaluations. To avoid bias from bad models only models with Z>0 are included in the global analysis. Single methods (blue), quasi (green), clustering (light grey)2 and combination models (dark grey). It is clear that using GDT_TS the consensus based methods are slightly better than the single-model predictors, while this is not the case using alternative measures. Clustering methods benefit a lot from having low quality models in the pool while the single model methods appear better at ranking higher quality models. For local correlation CAD values were not available so only the distances, turned into S-scores, and 1DDT values are compared. Here, for both measures the single evaluation methods are less good than the superposition based ones, but the difference is smaller when using 1DDT.

#### Comparison of global accuracy estimation predictions

How similar are the different model accuracy estimation scores produced by the different methods? To answer this we calculated the correlation between predictions from all methods, see Figure 4, and clustered methods using the Weighted Pair Group Method Centroid (WPGMC) with the median correlation as linkage. It can be seen that all methods (except qSVMQA) that use some sort of consensus (quasi-single or consensus) are clustered. Within this group the separation is primarily not between quasi-single methods and consensus methods, but rather between the methods that primarily use consensus and those who combine the consensus score with ProQ2. Pcomb-domain, ModFOLD6_rank, Wallner, and ModFOLD6 all use ProQ2 as part of their scoring and they all cluster together, while ModFOLD6_cor is more similar to the pure consensus methods (Pcons and ModFOLDclust2) than the other combined methods as it does not use ProQ2 global scores directly in its classification. Since the combined methods include single methods they are also more similar to all the single methods than the pure consensus methods.

**Figure 4:**
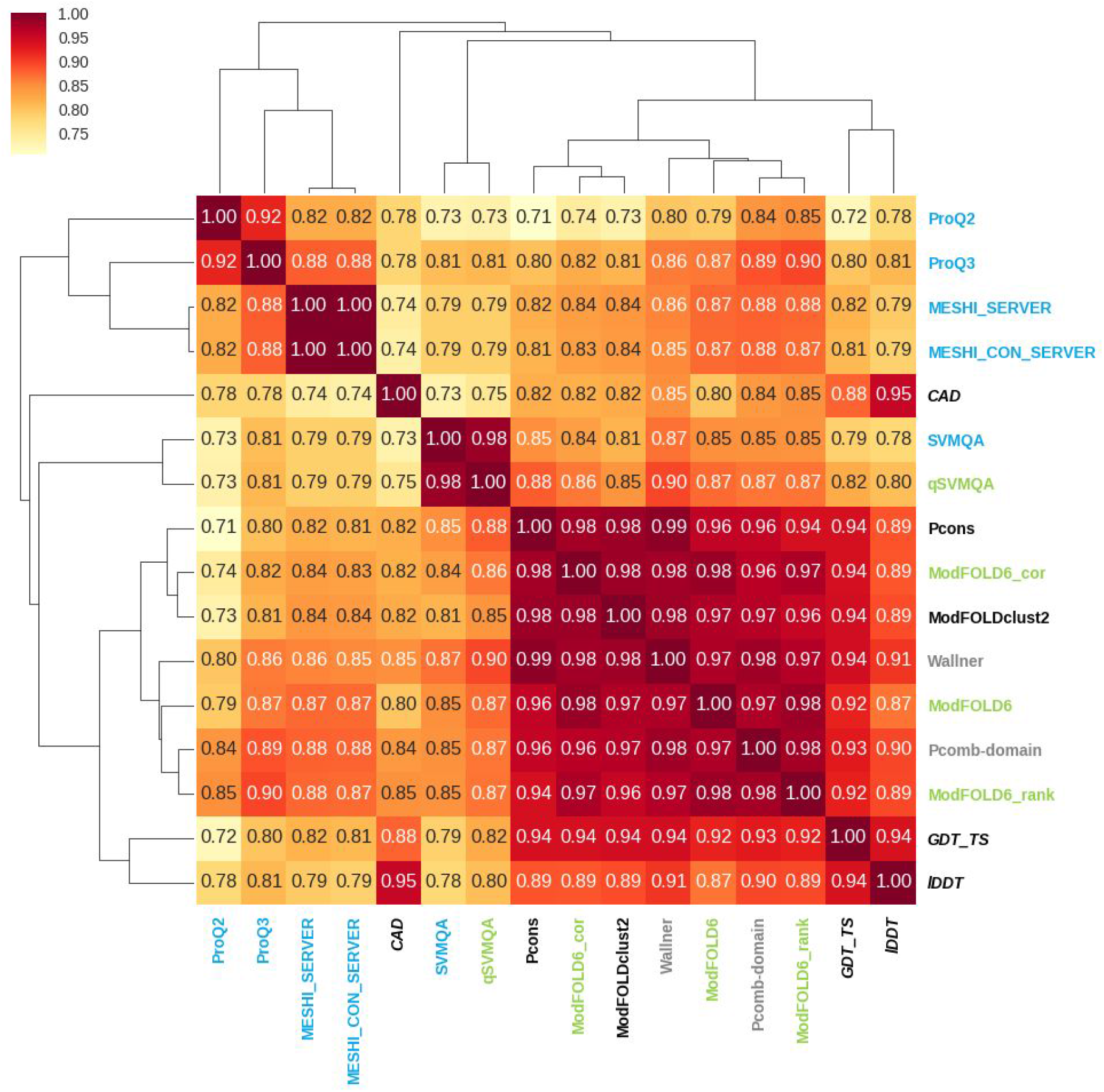
Pairwise correlations between predicted global accuracy scores from different methods and actual accuracy scores according to three measures. The methods are clustered hierarchically using WPGMC algorithm with the median correlation as similarity measure. Methods are colored as follows. Dark grey - pure consensus methods, light grey - combined single/consensus methods, green - quasi-single methods and blue pure single methods. It can be noted that (i) both quasi, pure and combined consensus methods are very similar (cc>0.94), while the single model quality methods are more different (cc<0.90 between the groups). ProQ2 is the real outlier only having a cc>0.82 to ProQ3 and the consensus methods that uses ProQ2 as a part of their score. In fact ProQ2 and ProQ3 are less similar to each other than any pair of consensus based methods. It can also be noted that the combined methods are more similar to the single-model methods than the pure consensus methods (Pcons, ModFOLDClust2).

Single model accuracy estimation methods show the largest performance diversity. SVMQA is the least similar to the others, being more similar to the consensus methods than to any other single model accuracy estimation method. The other three methods are more correlated, with the newer methods ProQ3 and MESHI showing the highest correlation. It can also be noted that in general ProQ2 is the outlier, showing the lowest correlation with the consensus methods.

When comparing the three different quality measurements (GDT_TS, CAD and 1DDT) it can be seen that they do not correlate with each other better than the correlation between the predicted values from the consensus methods and GDT_TS. The correlation between the quality measures of CAD and GDT_TS is 0.88 while the correlation of the predicted values from the consensus methods show a correlation to the GDT_TS values of 0.92 or higher, see Figure 4. As mentioned above some of the problems might origin from domain division, but it is clear that the model quality estimation accuracy is rivaling the ability to accurately measure the true quality of a model.

### Local accuracy estimation in CASP12

In terms of estimation of local accuracy, the best performance is obtained by the pure consensus methods followed by quasi-single model approaches^49^. In Figure 5 a heat map of the correlation between all local predictions by the methods discussed in this paper is shown. Unfortunately, of the single predictors evaluated here only ProQ2 and ProQ3 produce local predictions, nevertheless the trend is similar as for the global methods. All the consensus and quasi-single methods provide very similar accuracy estimates, while the two single model methods are outliers. It is clear from this analysis that the consensus methods correlate better with S-score (cc~0.85 vs ~0.65 for the ProQ methods) but less so with 1DDT (cc 0.77 vs. 0.71). As the consensus methods are based on a superposition algorithm this might not come as a surprise. Interestingly both ProQ2 and ProQ3 correlate better with 1DDT than with S-score. It can also be noted that ProQ3 correlates better than ProQ2 with both 1DDT and S-score, highlights the improvements made in single model quality estimates since CASP11.

**Figure 5:**
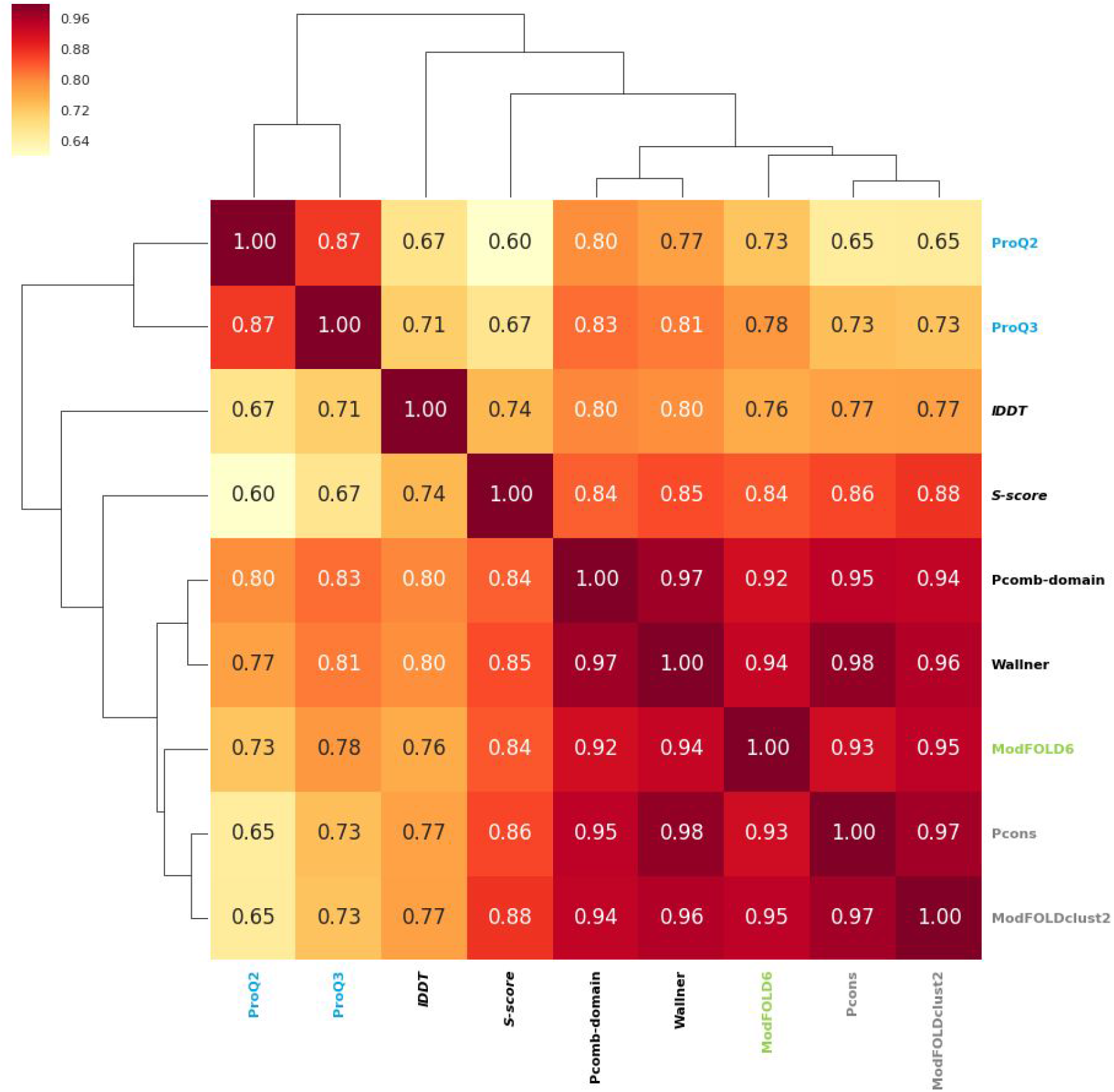
Pairwise correlation between local predicted S-scores calculated from the predicted distance using S-score formula (see above) with d_0_=5 and local 1DDT values (unfortunately local CAD scores were not available). Only methods that predicted local quality are included. As the ModFOLD6 methods only differ in their global scores and provide identical local estimates they were all represented by the ModFOLD6 method. Methods are colored as follows. Dark grey - pure consensus methods, light grey - combined single/consensus methods, green - quasi-single methods and blue pure single methods.

## Discussion

For the first time single model quality estimators can challenge the consensus based methods when it comes to ranking of targets. However, the consensus based estimators are still superior when it comes to local quality estimation, at least when using the CASP defined criteria. Below we will continue the CASP style of presentations by summarizing what each group learned during CASP12.

### What the Elofsson group learned

An interesting trend in CASP12 is that ProQ3 is better than our consensus method, Pcons, at picking up the best model (see EMA assessment paper^49^). In earlier CASPs this was not the case and until CASP10 it was clear that consensus based methods were superior even in this aspect. We do believe that the main reason for this is that single model accuracy estimation methods have improved in the last few years.

However, still consensus-based methods such as Pcons are superior at separating correct and incorrect models^49^. Interestingly, when using CAD, ProQ3 performs slightly better than Pcons even on this measure, see Figure 3, indicating that some part of the superior performance of consensus methods might be due to multi-domain properties of the targets or the choice of target function.

One issue at CASP is that the definition of the target function for local prediction used in CASP might not be ideal. The goal is to predict the error in distance for a particular residue. However, this is dependent on the superposition used, which can be problematic for multi-domain targets. It could therefore be useful in future CASPs to consider changing the target function to one of the non-superposition based quality evaluations, such as CAD or 1DDT. The stated goal in CASP12 is to predict the distance after superposition and for this consensus methods are better. However, the performance of ProQ3 is getting closer to the performance of the consensus methods when using 1DDT for model quality estimation, see Figure 3 and 5.

### What the Keasar group learned

The major rationale behind the design of MESHI-score pipeline (Figure 2) is to keep the feature set painlessly extendable. To this end we employed an ensemble-learning scheme, in which the feature selection is part of the training of each predictor (i.e. ensemble member). This way each feature has a “fair chance” to be included in some of the predictors and provide its unique contribution to the overall score. Overfitting at the single predictor level is avoided by restricting the number of selected features. Combining the set of predictor scores to form the single ensemble score (MESHI-score) does not require any adjustable parameters and thus, does not introduce overfitting at the ensemble level. In this experiment we put to test the modularity of our ensemble learning approach. Indeed, in this experiment we were able to get better results than before, simply by adding more features to the same machinery, with neither considerable computational burden nor overfitting. This encourages us to work on the development and adoption of more informative features.

In CASP12 we also tested MESHI-score-con for the first time, and its performance was a bit superior to that of MESHI-score. We take this as a proof of concept and wish to extend it in two directions: have a data-driven less restricted definition of the neighbors set, and apply the same idea also to decoys of high score. High scores to two dissimilar decoys must imply that at least one of them (often both) is wrong.

### What the Lee group learned

According to the CASP12 assessment, SVMQA is one of the best methods for selecting good quality models from a set of given decoys in terms of GDT-LOSS. The newly implemented features (five potential energy-based terms and consistency-based terms^45^) a systematic benchmarking approach on the selection of the final set of features, the optimization of machine learning parameters on a balanced training and testing dataset, and the usage of two separate predictors made SVMQA to perform significantly better than our old method used in CASP11 (RFMQA) when benchmarked on CASP11 targets. Additionally, SVMQA made valuable contribution to our tertiary structure prediction server (GOAL) and human predictors (LEE and LEEab) of CASP12 in terms of model selection. In terms of the model selection, SVMQA performed well, however, in term of assigning proper absolute global accuracy value to a model it didn’t perform as desired^49^. We believe that one way to improve on estimating the absolute score of a given model is to consider other types of objective functions to train separately for absolute global accuracy, which is one of the goals that we should work on for the next CASP.

### What the McGuffin group learned

The ModFOLD6 series of methods (ModFOLD6, ModFOLD6_rank and ModFOLD6_cor) perform particularly well in terms of assigning absolute global accuracy values. As expected the ModFOLD6_cor variant is the best of these as it was optimised for this task. The ModFOLD6 series of methods also perform competitively with clustering approaches for differentiating between good and bad models; the ModFOLD6_rank method being the best of these, which is only outperformed by two clustering groups (Wallner and Pcomb-domain). Furthermore, as we anticipated, the ModFOLD6_rank variant is better at selecting the top models than the ModFOLD6 and ModFOLD6_cor variants, however it is outperformed by the latest pure-single model methods. Overall, in terms of global scores, the ModFOLD6 variants rank within the top three methods for nearly every global benchmark according to 1DDT and CAD scores, as well as ranking within the top 10 according to other scores.

It is gratifying to see progress in CASP12 from many groups in both pure-single and quasi-single model approaches to estimate model accuracy. However, it is also clear there is still room for improvement of our methods. For instance, we are outperformed in terms of model selection by the newer pure single model methods. Further integration of methods is probably needed. Different methods are clearly better suited for different aspects of model accuracy estimation, therefore all approaches to the problem are still important to pursue. Perhaps the most difficult problem faced by all groups is how to optimize a global score for all aspects of model accuracy estimation, as there seems to be no one-size-fits-all solution presently. One potential solution to this might be to use a deep learning approach that outputs multiple scores depending on the intended use case. A global score for ranking models on a per-target basis, irrespective of the observed model-target similarity scores, is clearly very useful, if it can consistently select the better models. On the other hand a global score that can produce a near 1:1 mapping between predicted and observed scores, that is consistent across all targets, will allow us to assign accurate confidence scores to individual models (which is arguably more useful to an experimentalist than a top ranked, but nevertheless poor quality, model). Of course, as model accuracy estimation methods continue to improve and approach perfect optimisation for each use case, eventually the scores will converge on a single answer.

### What the Wallner group learned

Wallner and Pcomb-domain were the top two best method for differentiating between good and bad models (see assessment paper^49^). We were disappointed with the performance of Pcomb-domain, since in our benchmarks before CASP it would perform significantly better than Wallner. However, the true advantage of Pcomb-domain can only be seen if the assessment is performed based on domains or using superposition independent evaluation measures like 1DDT^52^ and CAD-score^51,52^. We calculated the per residue correlation of local predicted S-scores transformed using S-score formula (see above) based on either full-length target or target domains (Figure 6). For full-length assessment, methods based on global structural superposition (Wallner, Pcons, and ModFOLDclust2) for single domains are indeed superior. Also the performance based on multi-domain targets seems to be better for these methods (Figure 6a). However, the reason for this seemingly good performance for multi-domain targets is an artifact of the full-length assessment on multi-domain proteins that will only superimpose on one domain, if the domain-domain orientation is wrong. In effect, assigning high quality scores to the residues from one domain (usually the larger), and relatively low quality scores to the residues from other domains. This effect accentuates the performance for prediction methods using global superposition, which will also predict high quality scores for one domain and low scores for the others. If instead performance is measured using the official CASP domain definitions, this artifact can be avoided, and then Pcomb-domain performs better for multi-domain targets, and better than other methods when it uses a correct domain prediction (Figure 6b). Unfortunately, correct domain prediction was only achieved for 6 out of 21 multi-domain targets. Still, it pinpoints that there should be clear room for improving Pcomb-domain by improving the domain prediction algorithm.

**Figure 6:**
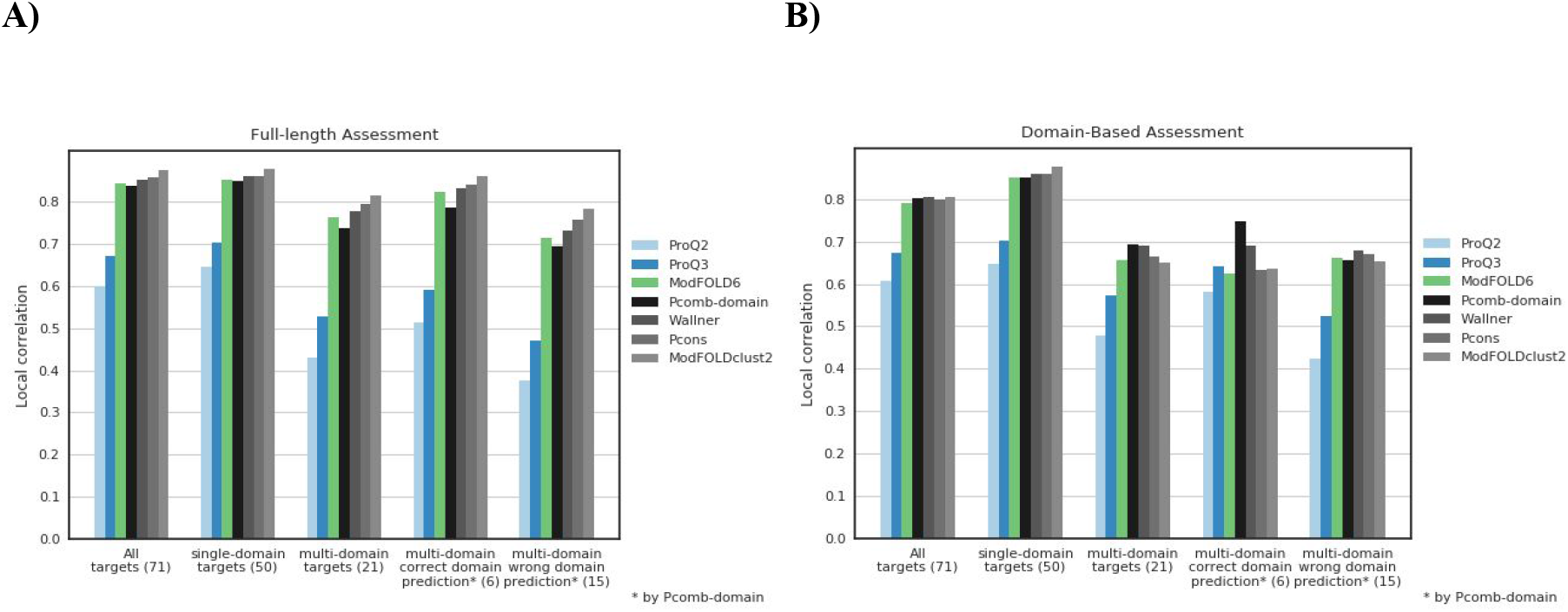
Per residue correlation of local predicted S-scores transformed using S-score formula with d_0_=5; based on full-length targets (A) and target domains (B) for selected methods and targets divided into multi and single domain targets. For full-length assessment methods based on superposition are superior. However, Pcomb_domain performs better than other methods when (and only when) it gets the domain prediction correct.

## Conclusions

It is our belief that the most important insight from the QA groups in CASP12 is the progress in single model accuracy estimations. Three new methods, SVMQA, MESHI and ProQ3, are all better than the best single model method in CASP11 (ProQ2). It is now clear that these methods are best at selecting the top-ranked model. However, quasi-single method and consensus methods are still superior when it comes to distinguishing correct and incorrect models and for local predictions. In those targets that have a wide spread of quality there is a clear distinction between the correlations of single and consensus methods with the later performing better. These are typically subunit of protein complexes, for which templates are available. Here, estimating the accuracy of a single model might not make sense without taking the entire complex into account. In CASP12 this is most dramatic with target T0865, where correlations for consensus based methods are high and correlations for all single model methods are negative. By comparing the predictions to each other it is seen that all consensus and quasi-single methods actually are very similar, while there is larger variation between the single methods, i.e. combining them might provide additional value in the future.

During this evaluation we noted issues for the multi-domain targets where predictors are successful at modeling the individual domains but not their relative arrangement. Here, the GDT_TS score (and any superposition based score) is based on the superposition of the largest domain. For CASP in general this has not been a problem as the evaluation is domain based, but for estimation of model accuracies the task is to evaluate the quality of an entire model and not of domains, as the domain division is not known at the time of predictions. This is most notably when evaluating local quality assessments. It could therefore be useful in future CASPs to consider changing the evaluation function for estimations of model accuracy to one of the non-superposition based quality evaluations, such as CAD^51^ or 1DDT^52^. Interestingly, when studying per target correlation single model estimations methods perform relatively better when assessed with CAD or 1DDT compared to using GDT_TS, see Figure 3 and 5. Probably for the same reason, the difference between consensus methods and single model estimation methods is smaller in domain-based than in full-length assessment, see Figure 6.

## Acknowledgements

First of all we are very grateful to all the work done by the late Prof. Anna Tramontano who has been fundamental for CASP. Her contribution will never be forgotten. We do also thank Dr. Andriy Kryshtafovych for his evaluation of our methods in CASP including providing additional data for our evaluations. We do also thank the rest of the CASP team for their efforts with CASP12. Finally we do acknowledge all the CASP participants who contributed with predictions that we could evaluate.

